# Differential requirement for the Ire1 luminal domain in *Candida albicans* drug susceptibility and pathogenicity

**DOI:** 10.64898/2026.09.24.753484

**Authors:** Samuel Stack-Couture, Nathan Towriss, Iwona Skulska, Anna Kalabina, Julie Genereaux, Vanessa Dumeaux, Rebecca S. Shapiro, Patrick Lajoie

## Abstract

The opportunistic human pathogen *Candida albicans* depends on the unfolded protein response (UPR) for cell wall integrity, antifungal tolerance, filamentous growth, and virulence. The UPR is driven by the conserved transmembrane sensor Ire1, which is activated either by misfolded proteins through its luminal domain or by lipid bilayer stress (LBS) through its transmembrane domain. In budding yeast, these two activation modes deploy divergent transcriptional programs. Whether the requirement for these two input domains is separable in *C. albicans*, where the cell membrane and cell wall are themselves the targets of major antifungal drug classes, remains unknown. Here, we engineered a *C. albicans* strain expressing Ire1 lacking an intact luminal domain (*ire1ΔLD*), which no longer effectively detects proteotoxic stress. The *ire1ΔLD* strain grew in the presence of the azole antifungals fluconazole and miconazole but was highly sensitive to heat shock, cell wall stress, and the echinocandin caspofungin. It was also unable to sustain filamentous growth and showed reduced virulence in a *Caenorhabditis elegans* infection model. RNA sequencing revealed only modest changes to the steady-state transcriptome of *ire1ΔLD* cells. Together, these findings define a differential requirement for the input domains of *C. albicans* Ire1, uncoupling growth under azole-induced membrane stress from the cell wall, thermal, and virulence-associated outputs that depend on proteotoxic sensing — a distinction that could inform antifungal strategies targeting the UPR.

## INTRODUCTION

The unfolded protein response (UPR) is a conserved transcriptional and post-transcriptional program that restores endoplasmic reticulum (ER) homeostasis when the organelle’s folding capacity is overwhelmed. In fungi, the UPR is driven by a single transmembrane sensor, Ire1, a kinase/endoribonuclease first identified in the budding yeast *Saccharomyces cerevisiae (Cox et al., 1993; Mori et al., 1993)*. Upon activation, *S. cerevisiae* Ire1 excises a non-spliceosomal intron from *HAC1* mRNA, and the cytoplasmic tRNA ligase Rlg1/Trl1 joins the exons to produce the active Hac1 transcription factor (Cox and Walter, 1996; Sidrauski and Walter, 1997; Sidrauski et al., 1996). Hac1 then drives a regulon of ∼380 genes encompassing ER chaperones, ER-associated degradation (ERAD) components, vesicular trafficking factors, and lipid-biosynthesis enzymes (Travers et al., 2000). In metazoans, an analogous *XBP1* splicing reaction performed by IRE1α extends this output to higher eukaryotes (Yoshida et al., 2001). The canonical model of UPR activation was derived from work in both yeast and mammalian systems and centers around the Ire1 luminal domain. Unfolded proteins in the ER lumen bind a peptide-binding groove on the Ire1 luminal domain and titrate BiP/Kar2 away from a regulatory site on the same domain. Together, this drives Ire1 oligomerization and RNase activation (Bertolotti et al., 2000; Credle et al., 2005; Gardner and Walter, 2011; Kimata et al., 2004).

Unfolded proteins are not the only manner by which Ire1 can be activated. Perturbations in ER membrane composition, collectively termed lipid bilayer stress (LBS), activate Ire1 through a separate mechanism that does not require the luminal domain (Fun and Thibault, 2020; Halbleib et al., 2017; Ho et al., 2020; Phuong et al., 2023; Pineau et al., 2009; Promlek et al., 2011). In *S. cerevisiae*, Ire1 variants lacking the core stress-sensing subregion of the luminal domain (ΔIII) or in which the luminal domain was replaced by a heterologous bZIP dimerization module remained responsive to inositol depletion and to *OPI3* deletion but failed to be acutely activated by tunicamycin or DTT, formally partitioning the two activation modes of Ire1 (Promlek et al., 2011). A variant lacking the entire luminal domain (*ire1ΔLD*) was subsequently shown to retain the ability to respond to LBS but was unresponsive to proteotoxic stress (Ho et al., 2020). This principle was then extended to mammalian cells, where it was shown that elevated levels of saturated phosphatidylcholine activates IRE1α and PERK through their transmembrane domains independently of misfolded proteins (Volmer et al., 2013). A study defining the molecular mechanism by which LBS activates Ire1 in *S. cerevisiae* showed that an amphipathic helix adjacent to the Ire1 transmembrane helix senses lipid bilayer compression and membrane packing defects, and that targeted mutations in this region selectively abolish LBS-induced activation while leaving the proteotoxic response intact (Halbleib et al., 2017).

The downstream consequences of LBS-driven Ire1 activation are not merely a milder version of the proteotoxic UPR. Using the *S. cerevisiae ire1ΔLD* strain coupled with genome-wide expression profiling, it was demonstrated that LBS and proteotoxic stress drive transcriptional programs that only partially overlap. The LBS-specific gene set was enriched for lipid metabolism and membrane biogenesis genes rather than the canonical chaperone and ERAD genes induced by tunicamycin (Ho et al., 2020). A parallel finding in *Caenorhabditis elegans* extended this principle to metazoans. LBS induced by phosphatidylcholine depletion upregulated a largely non-overlapping set of UPR target genes compared to tunicamycin treatment, including IRE-1–XBP-1-dependent activation of autophagy (Koh et al., 2018). The two stress inputs therefore not only converge on the same sensor through different domains, they also diverge downstream into distinct adaptive responses. This concept is supported by earlier *S. cerevisiae* observations that the UPR coordinates lipid biosynthesis and ER membrane expansion as homeostatic outputs (Cox et al., 1997; Schuck et al., 2009). Whether this transcriptional divergence is conserved beyond budding yeast and *C. elegans*, and in particular whether it operates in pathogenic fungi where the cell membrane and cell wall are the targets of major antifungal drug classes, has not been tested.

The opportunistic human pathogen *Candida albicans* offers a context where this question carries particular weight. *C. albicans* is the most common cause of invasive candidiasis, and its ability to switch between yeast and filamentous forms is crucial for invading host tissues, evading the immune system, and forming biofilms (Pappas et al., 2018; Shapiro et al., 2011; Soriano et al., 2023; Talapko et al., 2021). In *C. albicans*, Ire1 is required for filamentation, cell wall integrity, biofilm formation, antifungal tolerance, and virulence (Blankenship et al., 2010; Lee et al., 2023; Sircaik et al., 2021; Wimalasena et al., 2008; Zhao et al., 2025). Importantly, several of these functions are separable from Hac1. Ire1, but not Hac1, is required for plasma membrane localization of the high-affinity iron permease Ftr1 and for virulence (Ramírez-Zavala et al., 2022); Ire1 attenuates ER stress in part by activating the HOG-MAPK pathway in a manner that may not involve Hac1 (Husain et al., 2021); and Ire1 and Hac1 contribute differently to manganese tolerance (Henry et al., 2024). Although deletion of *HAC1* was initially reported to abolish filamentation (Wimalasena et al., 2008), *hac1Δ/Δ* cells are only partially impaired relative to *ire1Δ/Δ* (Lee et al., 2023), and an *IRE1*-depleted strain is sensitive to calcofluor white without undergoing *HAC1* splicing (Sircaik et al., 2021). We recently showed that the Ire1-dependent transcriptional program driving morphogenesis is largely distinct from the canonical proteotoxic UPR, that Ire1 but not Hac1 is required for upregulation of GPI-anchored cell wall genes during filamentation, and that Ire1 drives Hac1-independent decreases in transcripts encoding secretory proteins (Stack-Couture et al., 2026). These findings indicate that *C. albicans* Ire1 has acquired or retained outputs beyond *HAC1* splicing, but they do not address whether the two Ire1 input domains are differentially required for the drug tolerance and virulence outputs of the sensor protein.

To address this gap, we generated a *C. albicans* strain expressing Ire1 with a truncated luminal domain (*IRE1ΔLD*), adopting the genetic strategy that established the LBS-specific UPR in *S. cerevisiae (Ho et al., 2020; Promlek et al., 2011)*. We assessed the ability of *C. albicans ire1ΔLD* to support growth under proteotoxic, cell wall, cell membrane, and heat-shock stresses, and to undergo filamentation, and to establish virulence in a *C. elegans* infection model. We then used RNA-seq to define the steady-state transcriptome of cells in which Ire1 signals only through its transmembrane region, in the absence of an intact luminal domain. The results define a differential requirement for the two input domains of *C. albicans* Ire1: the luminal, proteotoxic-sensing arm is dispensable for growth under azole-induced membrane stress but required for cell wall stress tolerance, thermotolerance, filamentation, and full virulence.

## RESULTS AND DISCUSSION

### Differential requirement for the Ire1 luminal domain across canonical ER stressors and antifungal drugs

To study LBS-specific activation of Ire1 in *C. albicans*, we first constructed a strain expressing Ire1 lacking a major portion of the luminal domain (LD; Fig. 1A). Whereas the wild-type *IRE1* open reading frame contains a 1584 bp region encoding the luminal domain (codons 1–528, ending immediately upstream of the juxtamembrane amphipathic helix), our *IRE1ΔLD* construct removes 993 bp from within this region, deleting amino acids 78–408 in frame while retaining the signal sequence (amino acids 1–25) and the remaining 197 luminal residues. This *IRE1ΔLD* plasmid was transformed into the *ire1DX* strain, which has diminished expression of *IRE1* caused by deletion of one *IRE1* allele and replacement of the other allele’s promoter with the weakly expressed *PGA5* promoter (Woolford et al., 2016). The resulting *ire1DX+IRE1-ΔLD* strain is hereafter referred to as *ire1ΔLD*. In parallel, *ire1DX* was complemented with wild-type *IRE1* to generate the *ire1DX+IRE1-WT* strain, which served as the control throughout this study.

**Figure 1.**
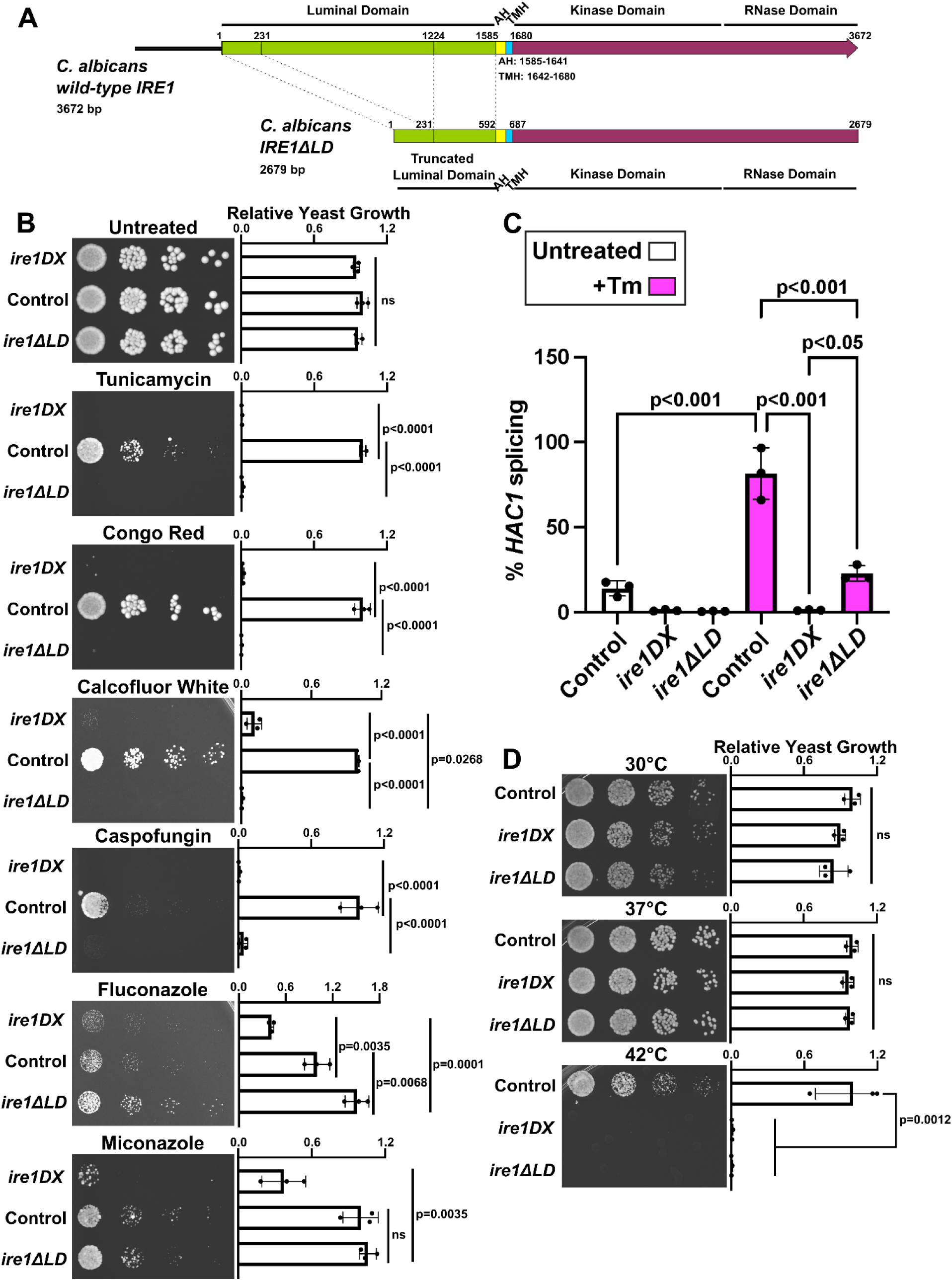
**The Ire1 luminal domain is differentially required for *C. albicans* growth in the presence of proteotoxic stressors, cell wall stressors, antifungal drugs, and heat shock.** A. Schematic of the *C. albicans* wild-type *IRE1* open reading frame (3672 bp) and the *IRE1ΔLD* construct (2679 bp), drawn to scale with nucleotide coordinates. Dashed lines indicate the boundaries of the deleted region. AH = amphipathic helix (bp 1585–1641), TMH = transmembrane helix (bp 1642–1680) (Halbleib et al., 2017), LD = luminal domain. **B.** Control (*ire1DX+IRE1-WT*), *ire1DX* and *ire1ΔLD* cells were spotted on YPD (untreated) or on plates containing 1.5 μg/mL tunicamycin, 10 μg/mL Congo red, or 10 μg/mL calcofluor white, 0.04 μg/mL caspofungin, 0.7 μg/mL fluconazole, or 0.03 μg/mL miconazole and incubated at 30°C. Calcofluor white images were acquired after 30 hours; all others after 24 hours. Growth was quantified by densitometry and normalized to the control strain on the same plate (Relative Yeast Growth, right panels). n=3, mean±SD plotted; one-way ANOVA with Tukey’s multiple comparison test. Exact p values are shown; ns, not significant. C. *HAC1* splicing was assessed in the indicated strains by RT-qPCR using primers amplifying either total or spliced *HAC1*, in untreated cells (white bars) or after treatment with 1.5 μg/mL tunicamycin for 2 hours (+Tm, magenta bars). For each strain, splicing is expressed as the percentage of spliced relative to total *HAC1*, calculated from 2^−ΔΔCt^ values. n=3, mean±SD shown; one-way ANOVA with Tukey’s multiple comparison test. Exact p values are shown. **D.** Strains were spotted on YPD plates and incubated at 30°C, 37°C or 42°C; images were acquired after 24 hours. Quantification and statistical analysis as in B.

As expected, *ire1*ΔLD could not grow on agar plates containing tunicamycin (Fig. 1B). Consistent with a compromised proteotoxic-sensing arm, *HAC1* splicing in *ire1*ΔLD was strongly attenuated following tunicamycin treatment, reaching only ∼23% compared with ∼81% in the control, and basal splicing was absent (Fig. 1C). Residual splicing in a luminal-domain mutant is not unexpected: in *S. cerevisiae*, the ΔIII mutant, which lacks the subregion of the core stress-sensing region required for unfolded-protein binding, loses acute activation by DTT or tunicamycin but gradually accumulates spliced *HAC1* (Promlek et al., 2011). Because our construct removes amino acids 78–408 while retaining the remaining luminal residues, it is best regarded as a large internal deletion of the stress-sensing region rather than a complete replacement of the luminal domain as used in *S. cerevisiae* (Ho et al., 2020), and the partial retention of splicing capacity is consistent with that design. Importantly, the residual splicing observed here is insufficient to support growth on tunicamycin, indicating that a threshold of Ire1 output, rather than any splicing at all, is required to survive acute proteotoxic stress.

*ire1*ΔLD was also unable to grow on plates containing the cell wall stressors Congo red and calcofluor white (Fig. 1B), which was expected given that cell wall integrity is closely tied to Ire1 and that cell wall stress leads to UPR activation (Blankenship et al., 2010; Scrimale et al., 2009). We also assessed the involvement of the Ire1 luminal domain in tolerating antifungal drug treatment. Caspofungin is an antifungal of the echinocandin class, which kills yeast by inhibiting the synthesis of β-(1,3)-D-glucan, a key component of the fungal cell wall (Sucher et al., 2009; Szymański et al., 2022). Consistent with growth on the other cell-wall-stressing agents, *ire1ΔLD* could not grow on plates containing caspofungin (Fig. 1B). However, *ire1ΔLD* was able to grow on plates containing fluconazole and miconazole, two azole antifungals (Fig. 1B). Azoles arrest yeast growth by inhibiting ergosterol synthesis in the cell membrane (Hitchcock et al., 1990); it is therefore likely that the absence of proteotoxic stress upon azole treatment permits *ire1ΔLD* to grow.

Interestingly, we found that loss of an intact Ire1 luminal domain rendered *C. albicans* cells acutely sensitive to growth at 42°C (Fig. 1D), pointing to a strong dependence on proteotoxic-stress sensing by Ire1 for survival at elevated temperature. This contrasts with *S. cerevisiae*, where Ire1 makes only a modest contribution to growth at elevated temperature: culturing at the maximum growth temperature of 39°C induces a weak (∼2-fold) increase in *HAC1* splicing, and *ire1Δ* cells grow only somewhat slower than *IRE1*+ cells under this condition (Hata et al., 2022). Notably, this heat-induced UPR was attributed to ER accumulation of unfolded proteins rather than lipid bilayer stress, as it was abolished in a luminal-domain mutant (ΔIII) but retained in a transmembrane mutant (V535R) (Hata et al., 2022). Although the temperatures are not directly comparable, as 42°C exceeds the near-maximal 39°C used in *S. cerevisiae* and falls within a host-relevant febrile range tolerated by *C. albicans*, both findings implicate the luminal, proteotoxic-sensing arm of Ire1 in the heat response. The far greater dependence of *C. albicans* on this arm for survival at elevated temperature may reflect its lifestyle as a warm-blooded-host pathogen, for which robust thermal adaptation is essential to virulence (Leach et al., 2012). The heat-shock growth defect of *ire1ΔLD* was also partially rescued by osmotic support with 1 M sorbitol (Supp. Fig. 1A,B), suggesting that loss of the luminal domain compromises membrane and cell wall integrity at elevated temperature rather than abolishing thermotolerance outright. To determine whether this thermal sensitivity reflected a broader failure to mount a heat-stress response, we examined two heat-inducible genes by RT-qPCR. Both *HSP104* and *ITR1* have previously been identified as direct targets of the heat shock transcription factor Hsf1 — the master regulator of thermal adaptation in *C. albicans* — with Hsf1 binding their promoters constitutively and in a heat-induced manner, respectively (Leach et al., 2016). *ITR1* and *HSP104* were robustly and comparably induced at 42°C across all strains, including ire1ΔLD (Supp. Fig. 1C,D), indicating that the heat sensitivity of *ire1ΔLD* does not arise from a failure to activate the canonical Hsf1-driven heat shock program. Rather, these data point to a specific requirement for the Ire1 luminal, proteotoxic-sensing arm in surviving elevated temperature, consistent with the loss of *HAC1* splicing and the heat dependence attributed to the luminal domain in *S. cerevisiae (Hata et al., 2022)*.

Surprisingly, *ire1ΔLD* and *ire1DX* were both able to grow on agar plates lacking inositol (Supp. Fig. 2A). Despite the ability of inositol depletion to induce the UPR through *HAC1* splicing (Sircaik et al., 2021), this contrasts sharply with *S. cerevisiae*, where UPR signaling is required to derepress *INO1* during inositol starvation (Brickner and Walter, 2004), such that *ire1Δ* and *hac1Δ* mutants are classically inositol auxotrophs (Cox et al., 1993; Nikawa and Yamashita, 1992). The ability of both *ire1ΔLD* and *ire1DX* to grow without exogenous inositol therefore indicates that, unlike in budding yeast, *C. albicans* does not depend on an intact Ire1 (or on its luminal, proteotoxic-sensing arm) to sustain *de novo* inositol biosynthesis. To ask whether inositol biosynthesis is induced normally in these strains, we measured *INO1* expression by RT-qPCR. Inositol withdrawal induced *INO1* by more than two orders of magnitude in wild-type and *ire1ΔLD* cells alike (Supp. Fig. 2B), demonstrating that neither an intact Ire1 luminal domain nor full *IRE1* expression is required to derepress *INO1* in *C. albicans*. Notably, *ire1DX* cells showed an approximately three-fold higher basal *INO1* level in the presence of inositol and a correspondingly blunted induction upon withdrawal, suggesting that diminished Ire1 expression partially derepresses the gene under inositol-replete conditions. This is consistent with a broader transcriptional rewiring of the *C. albicans* inositol regulon: the canonical *S. cerevisiae* regulators of Sc*INO1* — the repressor Opi1 and the Ino2–Ino4 activator heterodimer — do not control *INO1* in *C. albicans* (where they are instead repurposed to regulate, respectively, *SAP2*/morphogenesis and ribosomal protein genes), even though *INO1* remains responsive to extracellular inositol through as-yet-unidentified factors (Chen et al., 2008; Chen et al., 2015). Together these findings uncouple Ire1 and the conserved inositol-regulon circuitry from inositol homeostasis in this pathogen.

A caveat of this study is that lipid bilayer stress is not readily assayed through *HAC1* splicing in *C. albicans*. In *S. cerevisiae*, inositol depletion and *OPI3* deletion activate Ire1 through the transmembrane region and provide a clean readout of LBS sensing (Ho et al., 2020; Promlek et al., 2011). The situation in *C. albicans* is less straightforward: inositol depletion induces *HAC1* splicing, whereas neither fluconazole nor calcofluor white does so even after five hours of exposure (Sircaik et al., 2021), indicating that membrane and cell wall perturbations are not uniformly routed through the splicing arm of Ire1. Indeed, in *S. cerevisiae*, pharmacological inhibition of ergosterol biosynthesis with fluconazole prevents Ire1 from assembling into the clusters associated with full UPR activation (Cohen et al., 2017), raising the possibility that azoles dampen rather than engage Ire1 signalling — a further reason that azole treatment is a poor probe of transmembrane-arm activation. Consistent with this, we find that *ire1ΔLD* and *ire1DX* remain inositol prototrophs and induce *INO1* normally upon inositol withdrawal (Supp. Fig. 2B), so inositol depletion does not report Ire1 activation status in this organism as it does in budding yeast. We therefore restrict our conclusions to the requirement for the luminal domain; establishing whether the *C. albicans* Ire1 transmembrane region can be activated by membrane perturbation will require an LBS stimulus validated in this species.

### Loss of the Ire1 luminal domain modestly perturbs the steady-state transcriptome

To characterize the *ire1ΔLD* strain, we performed RNA sequencing. *ire1ΔLD* and control (*ire1DX+IRE1-WT*) cells were grown untreated for four hours prior to RNA purification and subsequent sequencing. Dimension reduction by principal component analysis separated *ire1ΔLD* from control samples along PC1, with the biological replicates of each genotype grouping together (Supp. Fig. 3A). Analysis of differentially expressed genes revealed 62 upregulated and 16 downregulated genes in *ire1ΔLD* relative to the control (|fold change|>2, padj<0.05; Fig. 2A). The six most highly upregulated genes in *ire1ΔLD* comprised five membrane transporters — *ATO1*, *HGT2*, *HGT12*, *HGT17* and *JEN2* — and *CTN1*, a carnitine acetyltransferase involved in cellular respiration (Supp. Table 2). The *HGT* family of genes is involved in glucose transport (Fan et al., 2002), whereas *ATO1* and *JEN2* facilitate the transport of acetate and dicarboxylic acids, respectively (Alves et al., 2017; Alves et al., 2025; Danhof and Lorenz, 2015). In accordance with these functions, gene ontology analysis for biological processes highlighted transmembrane transport (GO:0055085; padj=7.69E-03) — specifically carbohydrate and organic hydroxy compound transport (GO:0008643, padj=1.18E-04; GO:0015850, padj=2.56E-02) — as significant terms in the upregulated genes (Fig. 2B). Similarly, gene ontology analysis for molecular functions in the upregulated genes identified terms for various transporter activities (Fig. 2B).

**Figure 2.**
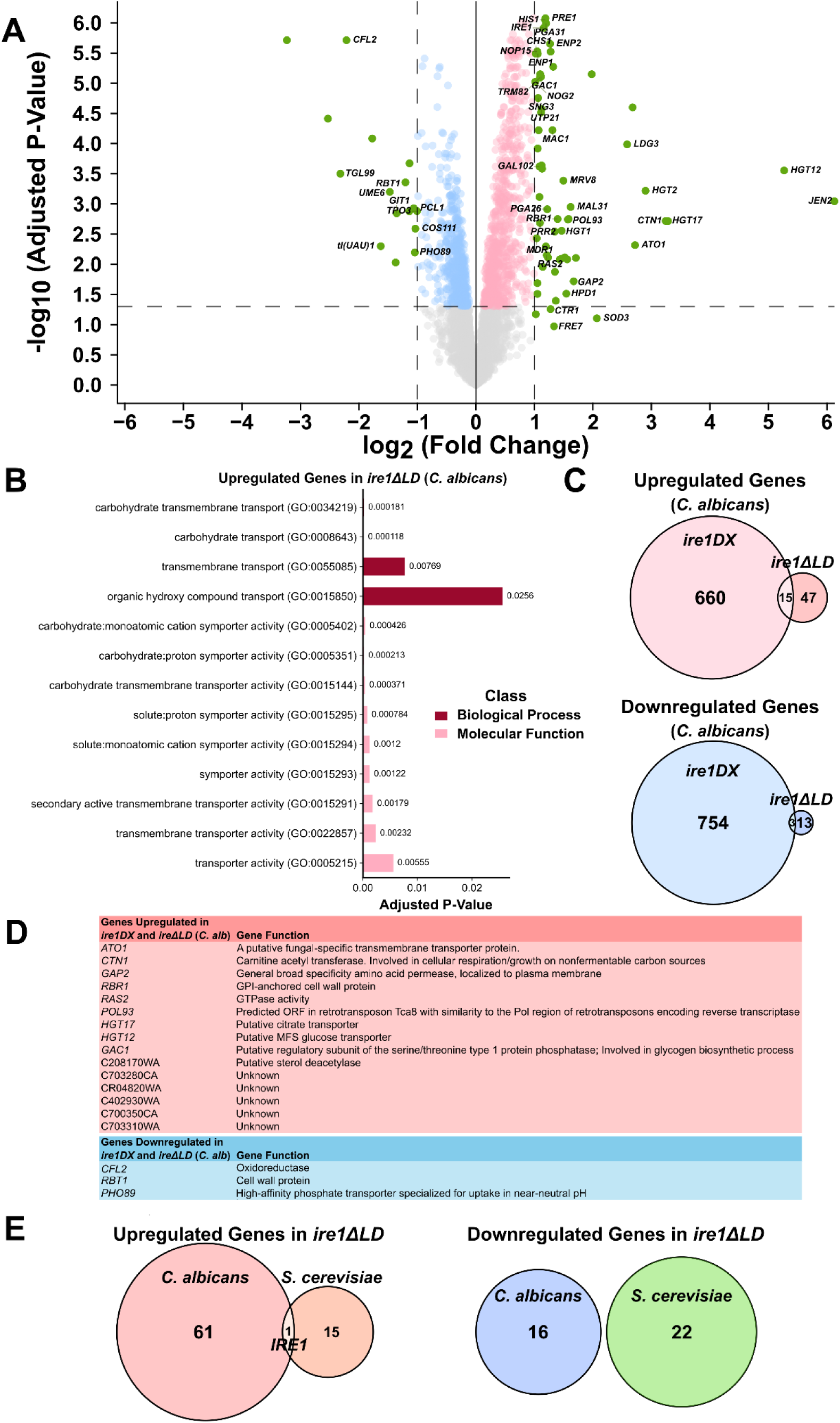
*C. albicans* cells lacking the Ire1 luminal domain have minimal changes to the steady-state transcriptome. **A.** Volcano plot showing differential gene expression. Genes highlighted in green have a |fold change|>2 (cutoff indicated by vertical lines). The horizontal dotted line represents a cutoff p_adj_ of 0.05. Only genes with a standard name are labeled. **B.** Gene ontology analysis for biological processes and molecular function in upregulated *C. albicans ire1ΔLD* genes (n=62). Analysis was performed using PANTHER via the Gene Ontology Consortium (Ashburner et al., 2000; Gene Ontology Consortium, 2026; Thomas et al., 2022). **C.** A comparison of genes up- and downregulated in *C. albicans ire1DX (Stack-Couture et al., 2026)* and *ire1ΔLD* relative to control strains. For both data sets, genes with |fold change|>2 and p_adj_<0.05 were considered for comparisons. **D.** Analysis of genes that are up- or downregulated in both *C. albicans ire1ΔLD* and *ire1DX* with |fold change|>2 and p_adj_<0.05. Gene function was obtained from Candida Genome Database (Lew-Smith et al., 2025). **E.** A comparison of genes up- and downregulated in *S. cerevisiae* and *C. albicans ire1ΔLD*. For *S. cerevisiae* (Ho et al., 2020), genes with |fold change|>1.5 and p<0.05 are considered; for *C. albicans*, genes with |fold change|>2 and p_adj_<0.05 are considered.

Of the 16 significantly downregulated genes in *ire1ΔLD*, nearly half encode proteins with uncharacterized functions, and the remainder encode proteins with various roles such as oxidoreduction (*CFL2*; (Baek et al., 2008)), polyamine transport (*TPO3*; (Gaur et al., 2008)), glycerophosphodiester transport (*GIT1*; (Bishop et al., 2011)), phosphate transport (*PHO89*; (Acosta-Zaldívar et al., 2024)), and cell wall organization (*RBT1*; (Bègue et al., 2025)). A notable downregulated gene was *UME6*, a key regulator of morphogenesis and biofilm formation in *C. albicans* (Supp. Table 2; (Banerjee et al., 2013; Carlisle et al., 2009)). Gene ontology analysis for biological processes and molecular function revealed no significant terms.

Previously, we performed RNA-seq on *ire1DX* (Stack-Couture et al., 2026). Analysis comparing untreated *ire1DX* and control strains identified 675 upregulated and 757 downregulated genes in *ire1DX* relative to the control (|fold change|>2, padj<0.05; Supp. Table 3). Here, we asked whether *ire1DX* and *ire1ΔLD* shared any differentially expressed genes and identified 15 genes upregulated in both. These genes encode proteins with a wide range of functions, including transporters (*ATO1*, *HGT12*, and *HGT17*), an amino acid permease (*GAP2*; (Kraidlova et al., 2016)), a GPI-anchored cell wall protein (*RBR1*; (Lotz et al., 2004)), a GTPase (*RAS2*; (Zhu et al., 2009)), and a serine/threonine protein phosphatase regulator (*GAC1*; (Miao et al., 2024); Fig. 2C,D). Three genes were downregulated in both strains: *CFL2*, *RBT1*, and *PHO89* (Fig. 2C,D). We also noted variable expression of *IRE1* itself. In *ire1DX*, *IRE1* was one of the most significantly downregulated genes, consistent with previous reports (Sircaik et al., 2021; Woolford et al., 2016), whereas in *ire1ΔLD* it was one of the most highly upregulated genes. This suggests that under steady-state conditions, *ire1ΔLD* may attempt to compensate for the lack of a fully functioning *IRE1* by upregulating its expression. The limited overlap of differentially expressed genes highlights largely distinct steady-state transcriptional profiles in *ire1DX* and *ire1ΔLD*. Whereas *ire1DX* shows extensive up- and downregulation of genes involved in a wide range of processes (Supp. Fig. 3B,C), ire1ΔLD shows modest changes in genes primarily involved in transport (Fig. 2B-D).

We also sought to compare differentially expressed genes in *C. albicans ire1ΔLD* (*Caire1ΔLD*) to those in an *S. cerevisiae ire1Δ+IRE1ΔLD* (*Scire1ΔLD*) strain at steady state (Ho et al., 2020). DNA microarray analysis of untreated *Scire1ΔLD (Ho et al., 2020)* identified 16 upregulated and 22 downregulated genes relative to the untreated control (|fold change|>1.5, p<0.05). Of the *Scire1ΔLD* upregulated genes, only one — *IRE1* — was also among the upregulated *Caire1ΔLD* genes, and no downregulated genes were shared (Fig. 2E). This limited overlap may in part be due to the lack of identifiable orthologs between the two species; only 45% of upregulated and 31% of downregulated genes in *Caire1ΔLD* could be matched to an orthologous gene in *S.* cerevisiae (Supp. Table 4). However, the fact that *IRE1* is the single shared gene highlights the compensatory mechanism initiated in the *ire1ΔLD* strains due to the expression of a truncated *IRE1*.

We performed gene ontology analysis for biological processes on the untreated *Scire1ΔLD* genes using SPELL, a search engine that compiles data from hundreds of different *S. cerevisiae* gene expression microarray experiments (Hibbs et al., 2007). Although none of the terms for upregulated genes in *Scire1ΔLD* matched those in *Caire1ΔLD*, the majority of upregulated *Scire1ΔLD* terms were related to carbohydrate metabolic processes, which aligns with the enrichment in carbohydrate transport seen in *Caire1ΔLD*. The downregulated *Scire1ΔLD* genes were enriched in biological processes primarily related to amino acid and small molecule biosynthesis (Supp. Fig. 3D,E).

Taken together, the *ire1ΔLD* steady-state signature is small and dominated by transmembrane transporters, with little overlap with either the expansive *ire1DX* profile or the *S. cerevisiae ire1ΔLD* dataset beyond the shared compensatory upregulation of *IRE1* itself — arguing that loss of the luminal domain reshapes the transcriptome in a species- and lesion-specific manner rather than recapitulating a generic UPR-deficient state. The downregulation of *UME6*, a master regulator of filamentation and biofilm formation, is the most functionally suggestive change. *UME6* is known to play a key role in regulating morphogenesis and pathogenicity, with *UME6* overexpression driving the yeast-to-hyphae transition, biofilm development, and tissue invasion in a model of oropharyngeal candidiasis (Banerjee et al., 2013; Carlisle et al., 2009). RNA-seq and ChIP analysis have shown that Ume6 forms complexes with the master morphogenesis and biofilm regulators Efg1 and Ndt80 that enable it to bind the promoters of biofilm genes and extensively regulate pathogenicity (Do et al., 2025). It is therefore possible that the decreased expression of *UME6* observed in steady-state *ire1ΔLD* cells provides a candidate link between the truncated receptor and the morphogenetic defects examined below.

### The Ire1 luminal domain is required for morphogenesis and virulence in *C. albicans*

Given the key role for Ire1 in morphogenesis and virulence (Blankenship et al., 2010; Lee et al., 2023; Sircaik et al., 2021; Stack-Couture et al., 2026; Wimalasena et al., 2008; Zhao et al., 2025) and the varying roles for the Ire1 luminal domain in antifungal tolerance, we investigated the requirement for the Ire1 luminal domain in *C. albicans* pathogenicity. Previously, it was shown that *ire1DX* is unable to undergo filamentation under various morphogenesis-inducing conditions (including 10% fetal bovine serum at 37°C and Spider medium; (Sircaik et al., 2021; Stack-Couture et al., 2026)). Here, we find that *ire1ΔLD* is also unable to filament in serum at 37°C (Fig. 3A,B; 0.68% filaments, compared with 29.27% for the control). We also investigated *ire1ΔLD* virulence using a *Caenorhabditis elegans* infection model. When grown on a plate of *C. albicans* cells, *C. elegans* worms will ingest the yeast and develop a gastrointestinal infection that ultimately kills the worm. After 72 hours, we observed the lowest survival probability for *C. elegans* infected with control *C. albicans* (52.46% alive) and the highest when infected with *ire1DX* (Fig. 3C; 72.63% alive). Worms infected with *ire1ΔLD* had an intermediate survival probability relative to worms infected with the other two strains (64.19% alive), suggesting only a partial role for the Ire1 luminal domain in regulating *C. albicans* virulence.

**Figure 3.**
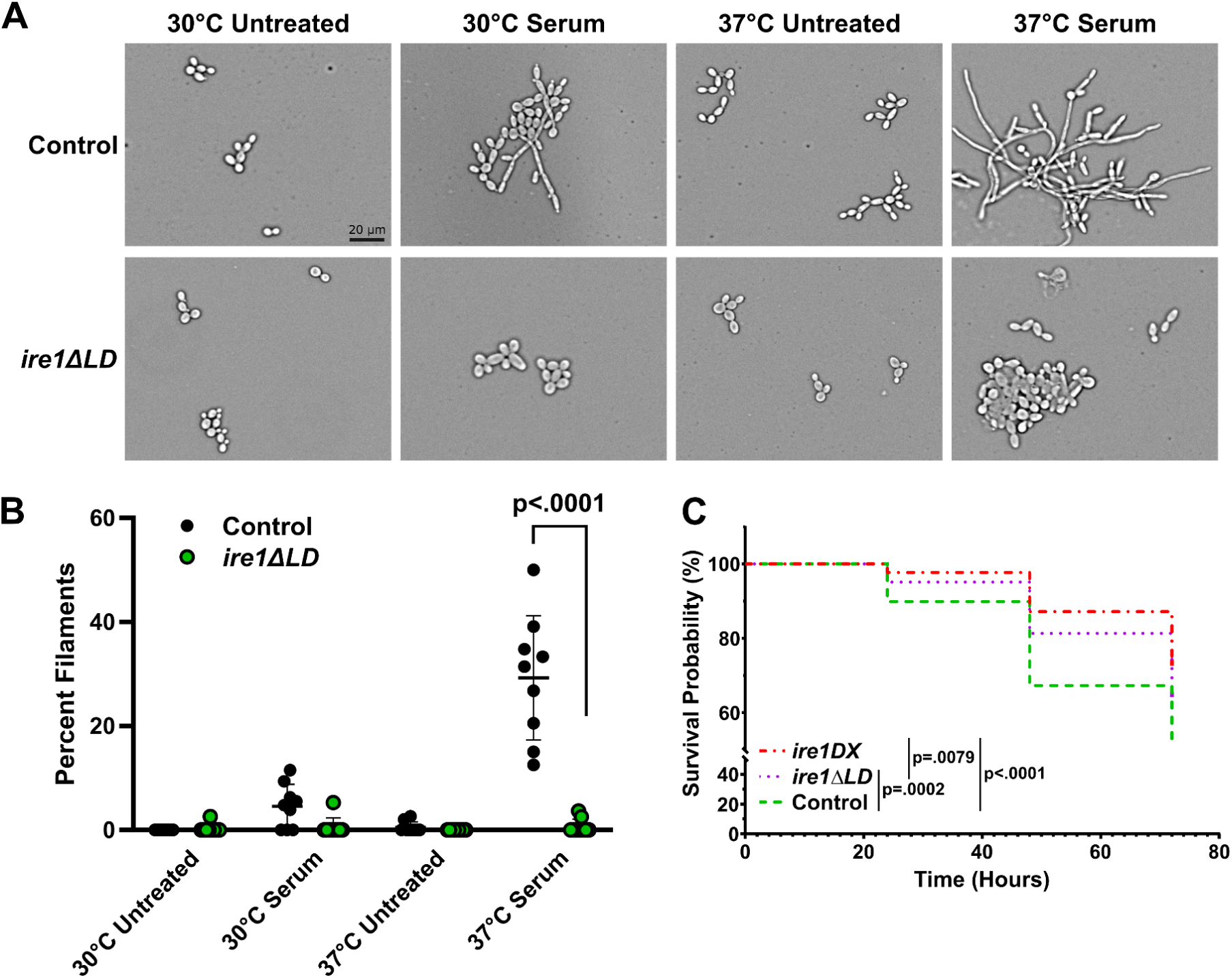
The Ire1 luminal domain is required for filamentous growth and virulence in *C. albicans*. **A.** The indicated *C. albicans* strains were grown in 10% fetal bovine serum for four hours at 37°C to induce filamentation or grown untreated at 30°C as a control. Images were obtained using the Cytation5 cell imaging multi-mode reader. Scale bar = 20 µm, applies to all images. **B.** Percent of cells in a filamentous form (Percent Filaments) was quantified for 3 biological replicates. Statistical analysis performed in Graphpad Prism using a one-way ANOVA with Tukey’s multiple comparison test, mean±SD plotted. **C.** Survival curve for AU37 *C. elegans* worms grown on plates containing a lawn of indicated *C. albicans* strains. Statistical analysis performed in Graphpad Prism using logrank (Mantel-Cox) test to compare conditions.

In summary, by removing the luminal domain of *C. albicans* Ire1, we define a differential requirement for its two input domains. The truncated receptor is sufficient to support growth under azole-induced membrane stress — including fluconazole and miconazole — whereas the luminal, proteotoxic-sensing arm is required for survival of cell wall stress, echinocandin treatment, and elevated temperature. The luminal domain is also needed for filamentous growth, and it contributes to virulence in a *C. elegans* infection model. Notably, this partitioning is accompanied by surprisingly modest steady-state transcriptional change, indicating that the luminal domain’s contribution is felt largely under stress rather than at baseline. Several of these behaviors diverge from *S. cerevisiae* — most strikingly the acute heat sensitivity of *ire1ΔLD* and the retention of inositol prototrophy — underscoring that Ire1 outputs have been rewired in this pathogen alongside the broader inositol regulon. Together, these findings show that membrane-stress adaptation can be separated from the pathogenic outputs of the full-length receptor, and they suggest that strategies disabling the proteotoxic-sensing arm of Ire1 could compromise *C. albicans* virulence and cell wall–targeted drug tolerance while leaving azole responses intact — a distinction worth weighing in UPR-directed antifungal design.

## MATERIALS AND METHODS

### Yeast Strains and Cell Cultures

The *C. albicans* strains used in this study are described in Supplemental Table 1. Strains were grown from frozen stocks at 30°C on YPD (1% yeast extract, 2% peptone, and 2% dextrose) or selective synthetic complete medium plates (SC with appropriate amino acids). All overnight liquid cultures were grown in a rotating drum incubator at 30°C in triplicate unless otherwise stated. SC medium lacking inositol was prepared using YNB with ammonium sulfate and without inositol (MP Biomedicals, Solon, OH, USA), and SC medium with inositol was prepared using regular YNB and was supplemented with an additional 100 μg/mL D-myo-inositol.

### Reagents

Stock solutions of tunicamycin (5 mg/mL in DMSO; Millipore Sigma, Burlington, MA, USA), fetal bovine serum (100%; Millipore Sigma), caspofungin (1 mg/mL in H2O; Millipore Sigma), D-myo-inositol (100 mg/mL in water; MP Biomedicals), sorbitol (2 M in water; Thermo Fisher Scientific, Waltham, MA, USA), Congo red (10 mg/mL in water; Thermo Fisher), fluconazole (10 mg/mL in DMSO; Thermo Fisher), miconazole (10 mg/mL in DMSO; Alfa Aesar, Haverhill, MA, USA) and calcofluor white (10 mg/mL in water; fluorescent brightener 28; Millipore Sigma) were prepared and used as described in the text.

### Strains and Plasmid Design

All plasmids and primers used in this study are listed in Supplemental Table 1. The *IRE1ΔLD* plasmid was constructed by cloning *C. albicans IRE1* lacking the codons for amino acids 78–408 (a 993 bp in-frame deletion within the luminal-domain-encoding region) into pDDB78-*IRE1WT* to replace the wild-type *IRE1*. The signal sequence was retained.

### Growth Assay on Agar Plates

Overnight cultures were diluted to an OD_600_ of 0.1 and serially diluted 1:5 in a 96-well plate. A 48-pin replica plater was used to spot equal volumes of cells onto agar plates. Plates were incubated at 30°C, or at 37°C or 42°C for heat shock experiments. Images were captured 20–30 hours after spotting using an SP Imager (S&P Robotics, North York, ON, Canada); see figure legends for specific information.

Quantification was performed using ImageJ (Schneider et al., 2012) following a protocol that measures the mean grey value of the spots on each plate (Petropavlovskiy et al., 2020). Briefly, images were converted to 8-bit format and background noise was removed using the rolling ball algorithm. The grey value of a spot from each condition was measured, using the same dilution within one experiment, and the mean grey value of the background was subtracted from this measurement. All values were normalized to the control strain on the same plate unless otherwise stated in the figure legend. Statistical analysis was performed in GraphPad Prism, version 11.0.1 (GraphPad Software, Boston, MA, USA).

### Filamentous Growth Assay

Filamentation assays were performed as previously described (Stack-Couture et al., 2026). Briefly, cell cultures were grown overnight and diluted to an OD_600_ of 0.2 in fresh medium. Cultures were grown to log phase for 1.5 hours, then split and either treated with 10% fetal bovine serum and incubated at 37°C or left untreated and incubated at 30°C for four hours. Samples were added to glass slides and imaged using a BioTek Cytation 5 Cell Imaging Multimode Reader. Three images were taken of each biological replicate from randomly selected regions of the slide. To quantify percent of cells in a filamentous form (Percent Filaments), round and filamentous yeast were counted from images by hand. Mean percent filaments was plotted ±SD for each condition. Statistical analysis was performed in GraphPad Prism.

### Infection Assay

Infection assays were performed as previously described (Razzaq et al., 2021). Briefly, *C. elegans* AU37 worms were bleached and resuspended to extract eggs and develop a fresh stock of worms. Worms were plated on superfood agar plates and incubated overnight at room temperature. Plates were then incubated for an additional two nights at 25°C. *C. albicans* strains were cultured in YPD overnight at 30°C with shaking, then spread onto BHI-kanamycin plates and incubated overnight at 30°C. The superfood agar plates were washed with S Basal to dislodge worms, and the worms were washed an additional three times with S Basal. Worms were then suspended in S Basal (2 worms/μL) and 80–100 worms were plated on the *C. albicans*-inoculated BHI-kanamycin plates. Plates were stored in a 25°C incubator for three days, and worms were scored each day by counting and removing them with a platinum wire worm pick under a stereomicroscope. Statistical analysis was performed in GraphPad Prism using the logrank (Mantel–Cox) test to compare conditions.

### RNA Purification

Control (*ire1DX+IRE1-WT*) and *ire1ΔLD* strains were cultured overnight in quadruplicate. In the morning, cells were diluted to an OD_600_ of 0.2 and grown for 90 minutes at 30°C to log phase. Cells were then grown for an additional four hours untreated. Total RNA was isolated using a MasterPure Yeast RNA Purification Kit (LGC Biosearch Technologies). DNase treatment was performed using a DNA-free DNA Removal Kit (Invitrogen). RiboGuard RNase inhibitor (1 µL) was added to each sample to prevent RNA degradation.

### RNA Sequencing

Quality control analysis of purified RNA samples was performed using the Agilent 4150 TapeStation (London Regional Genomics Centre, London, ON, Canada) to confirm an RNA integrity number of 7 or greater. Library preparation and RNA sequencing were performed by Plasmidsaurus using the Illumina platform with custom analysis and annotation. PolyA mRNA was captured and converted to double-stranded cDNA using reverse transcription and second-strand synthesis, followed by tagmentation, library indexing, and amplification. Differential gene expression was quantified using 3′ end counting. Each sample generated 12.7–14.5 million unique reads, with RNA yields of 39.07–45.98 ng/µL. FASTQ files and normalized counts were deposited in NCBI’s Gene Expression Omnibus (Edgar et al., 2002) and are accessible through GEO Series accession number GSE336334.

FASTQ files were generated using BCL Convert v4.3.6 and fqtk v0.3.1. Preprocessing and quality control were performed using fastp v0.24.0, which performed poly-X tail trimming and 3′ quality-based tail trimming using a Phred score of 15 (Chen et al., 2018). Any reads with a length below 50 bp were removed. Reads were aligned to the reference genome of *Candida albicans* SC5314 Assembly 22 (NCBI RefSeq assembly GCF_000182965.3; GenBank assembly GCA_000182965.3) using STAR aligner v2.7 with non-canonical splice junction removal and output of unmapped reads (Dobin et al., 2013). BAM files were sorted using samtools v1.21 (Danecek et al., 2021). PCR and optical duplicates were removed using UMICollapse v1.1.0 (Liu, 2019). Further quality control was performed with RustQC v0.2.1, and a quality control report was generated using MultiQC v1.33 (Ewels et al., 2016). featureCounts v2.1.1 was used for quantifying gene expression (Liao et al., 2014). Principal component analysis was calculated with counts normalized by the trimmed mean of M-values (Robinson and Oshlack, 2010) method using Pearson correlation (Schober et al., 2018). Differential expression analysis was performed using edgePython v0.2.5 (Chen et al., 2025). Gene ontology enrichment analysis was performed using PANTHER via the Gene Ontology Consortium (Ashburner et al., 2000; Gene Ontology Consortium, 2026; Thomas et al., 2022).

Comparison to *ire1DX* was performed using data accessible through GEO Series accession number GSE308581 (Stack-Couture et al., 2026). Gene functions were obtained from the Candida Genome Database (Lew-Smith et al., 2025). *S. cerevisiae* microarray data were accessed using GEO Series accession number GSE131146 (Ho et al., 2020). Gene ontology analysis for genes up- and downregulated in *S. cerevisiae* was performed using SPELL (Version 2.0.3r71) (Hibbs et al., 2007).

### RT-qPCR

Cell cultures were grown overnight in triplicate, then diluted 1:10 and grown to log phase for 90 minutes. Treatments were applied as described in figure legends. RNA was purified and treated with DNase as described in the RNA Purification methods. cDNA libraries were prepared using the SuperScript IV VILO Master Mix (Thermo Fisher Scientific) according to the manufacturer’s instructions, and RT-qPCR was performed using a SYBR Green Master Mix (Thermo Fisher Scientific) and a QuantStudio 3 Real-Time PCR System (Thermo Fisher Scientific). *ACT1* served as the housekeeping gene for all experiments; all RT-qPCR primers are listed in Supplemental Table 1. The comparative Ct method of analysis was used (Schmittgen and Livak, 2008), and unless otherwise stated, all ΔΔCt values were calculated by normalizing ΔCt values to the ΔCt of the untreated control strain. For *HAC1* splicing analysis by RT-qPCR, two sets of primers were used that amplify either total or spliced *HAC1*. The resulting 2^−ΔΔCt^ values were used to calculate a relative expression ratio of spliced to total *HAC1*. For *INO1* expression analysis, log-phase cells were washed and resuspended in synthetic complete medium either containing 100 μg/mL D-myo-inositol or lacking inositol, and grown for four additional hours at 30°C before RNA purification.

## ACKNOWLEDGEMENTS

PL is supported by an NSERC Discovery Grant (RGPIN-2022-05267) and CIHR Project Grants (PJT 168882 and ARB 192062). PL is also supported by a Western CIHR Accelerator grant as well as a Western CIHR Reapplication Program grant. VD is supported by an NSERC Discovery Grant (RGPIN-2024-04932). RSS holds the Canada Research Chair in Microbial Functional Genomics and Synthetic Biology. SSC was supported by an Ontario Graduate Scholarship.

