## Supplemental Figures for "Differential requirement for the Ire1 luminal domain in *Candida albicans* drug susceptibility and pathogenicity"

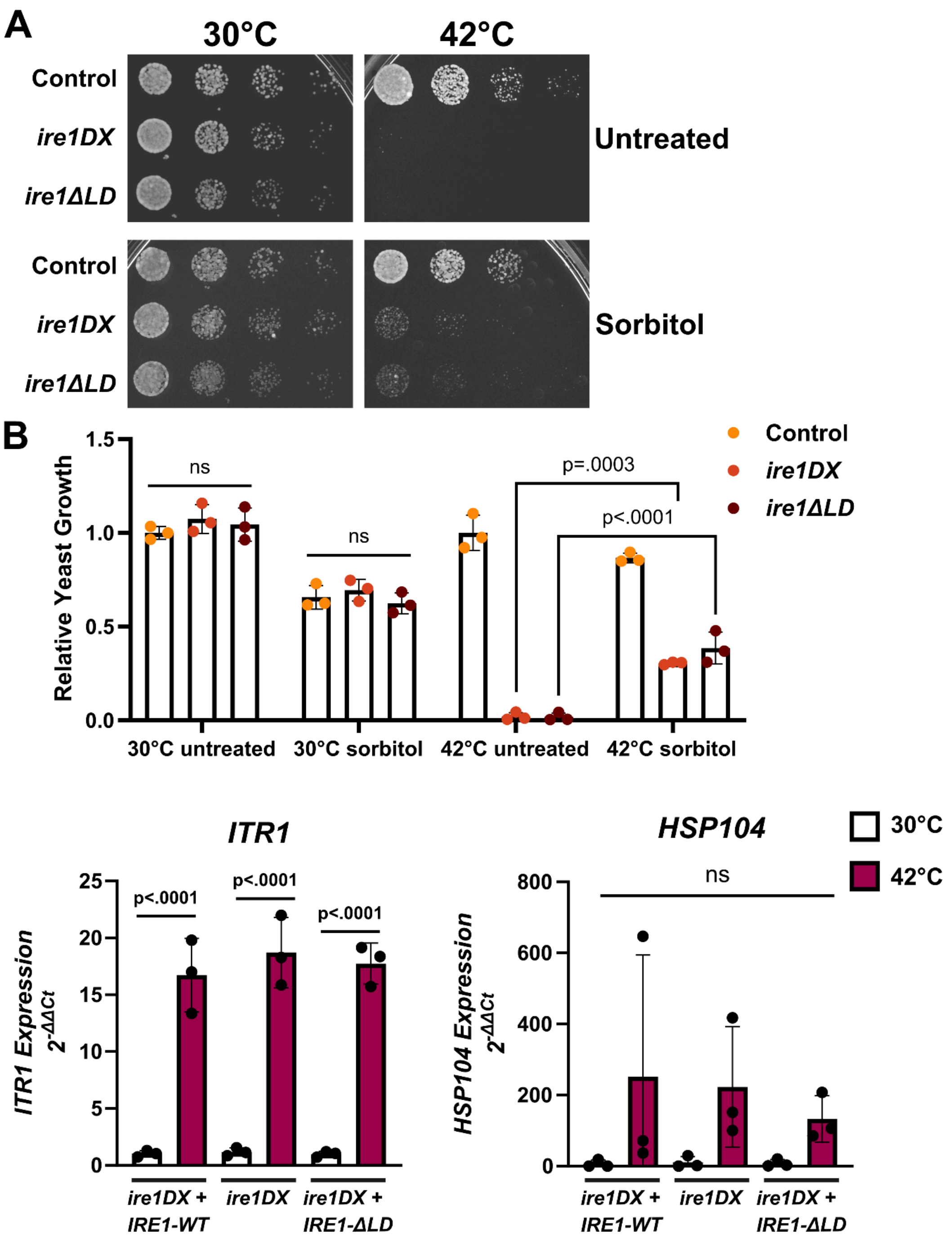


**Supplemental Figure 1. Growth of *IRE1*-deficient strains during heat shock can be rescued by sorbitol. A.** Indicated *C. albicans* strains were spotted on YPD plates with or without 1 M sorbitol and incubated at 30°C or 42°C for 24 hours. **B.** Cell growth was quantified and statistical analysis was performed using a One-way ANOVA with Tukey's multiple comparison. n=3, mean±SD plotted. Relative gene expression for **C.** *ITR1* and **D.** *HSP104* was assessed by RT-qPCR in indicated *C. albicans* strains grown at 30°C or 42°C for 60 minutes. Legend applies to both graphs. n=3, mean±SD shown, one-way ANOVA with Tukey's multiple comparisons performed in GraphPad Prism.

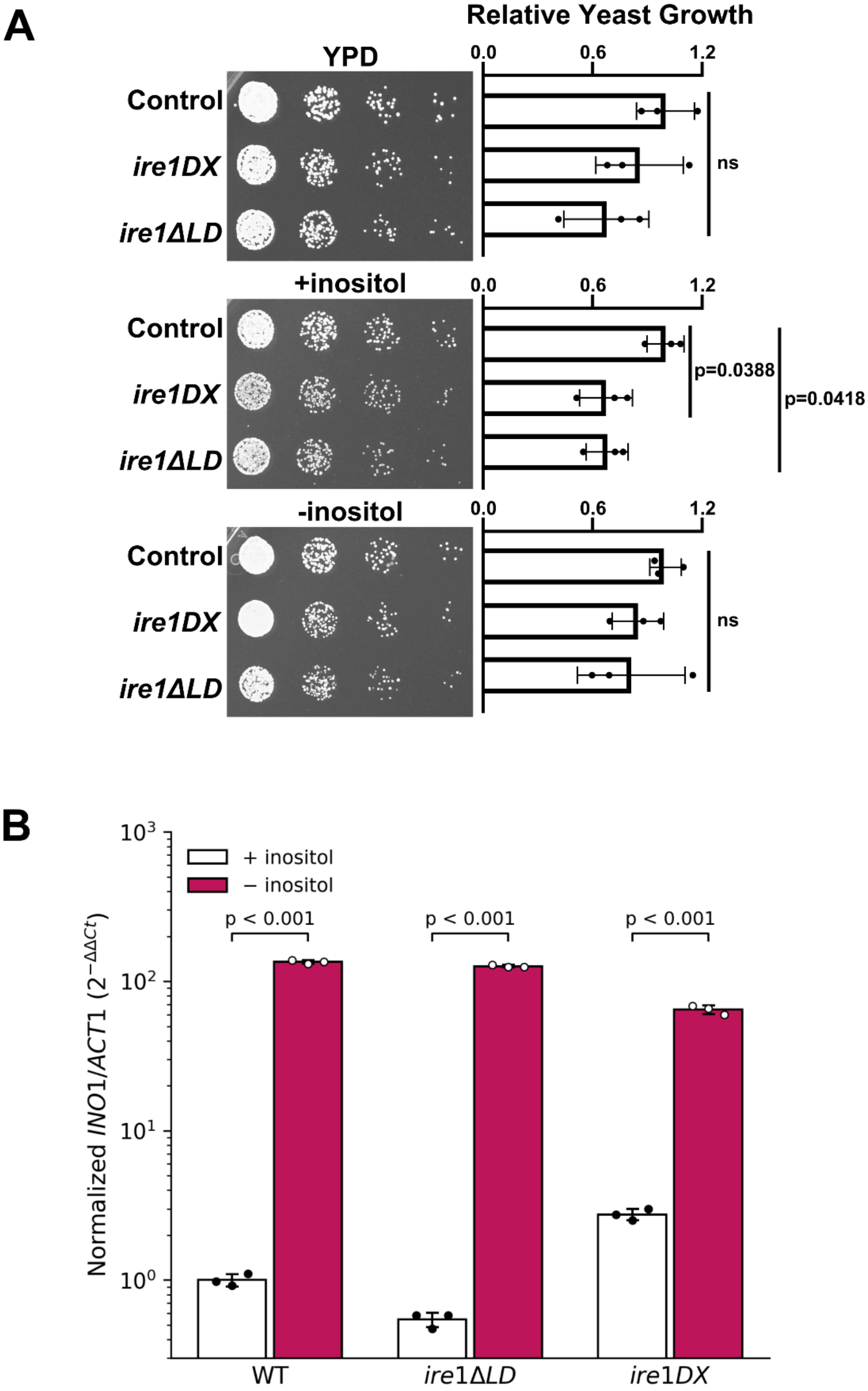
A
Relative Yeast Growth
YPD
0.0
0.6
1.2
Control
ire1DX
ire1ΔLD
ns
+inositol
p=0.0388
p=0.0418
-inositol
B
+ inositol
− inositol
p < 0.001
Normalized INO1/ACT1 (2^−ΔΔCt)
WT
ire1ΔLD
ire1DX

**Supplemental Figure 2. Deletion of the Ire1 luminal domain does not confer inositol auxotrophy in *C. albicans*. A)** *C. albicans* strains (control [*ire1DX+IRE1-WT*], *ire1DX*, and *ire1ΔLD*) were grown spotted on YPD, synthetic medium containing inositol (+ 100 µg/mL inositol), or synthetic medium lacking inositol (−inositol), and incubated at 30°C for 48 hours. Growth was quantified by densitometry and is expressed relative to the control strain on the corresponding medium (Relative Yeast Growth, right panels). n=3, mean ±SD plotted; statistical analysis was performed using a one-way ANOVA with Tukey's multiple comparisons in GraphPad Prism. ns, not significant. **B**) *INO1* mRNA levels were measured by qRT-qPCR in wild-type (WT), *ire1ΔLD* and *ire1DX* strains of *C. albicans* grown with (+ inositol, white bars) or without (− inositol for 4hours), magenta bars). *INO1* expression was normalized to *ACT1* and is shown as 2^−ΔΔCt relative to WT + inositol, which is set to 1.. Bars show mean ± SD of three biological replicates, each measured in technical quadruplicate. The y-axis is on a $\log_{10}$ scale. Two-way ANOVA (strain × inositol) on ΔCt values followed by Šídák's multiple comparisons test; $p < 0.001$ for + vs − inositol in each strain.

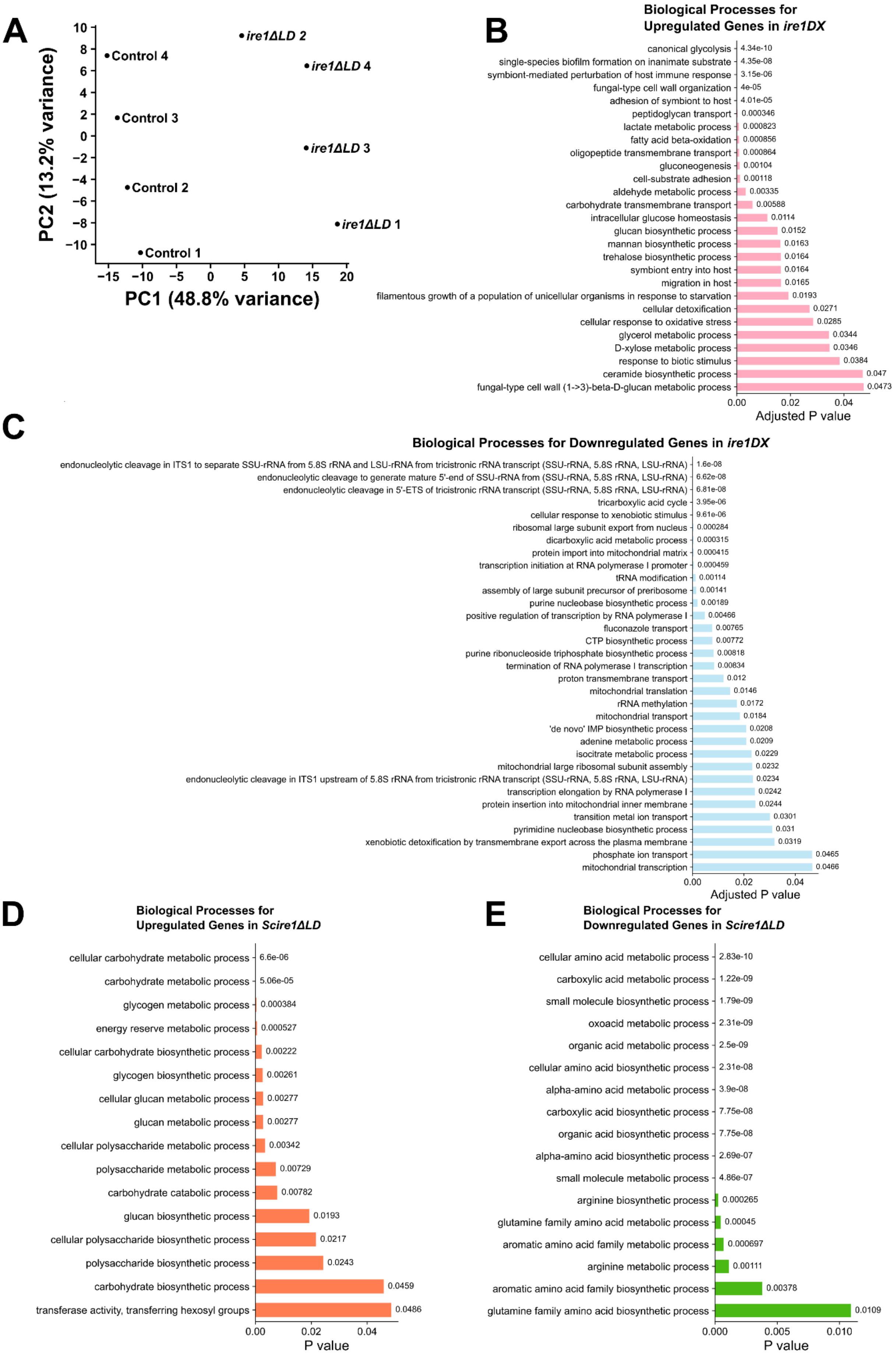
A
PC2 (13.2% variance)
PC1 (48.8% variance)
Control 4
Control 3
Control 2
Control 1
ire1ΔLD 2
ire1ΔLD 4
ire1ΔLD 3
ire1ΔLD 1
B
Biological Processes for Upregulated Genes in ire1DX
canonical glycolysis 4.34e-10
single-species biofilm formation on inanimate substrate 4.35e-08
symbiont-mediated perturbation of host immune response 3.15e-06
fungal-type cell wall organization 4e-05
adhesion of symbiont to host 4.01e-05
peptidoglycan transport 0.000346
lactate metabolic process 0.000823
fatty acid beta-oxidation 0.000856
oligopeptide transmembrane transport 0.000864
gluconeogenesis 0.00104
cell-substrate adhesion 0.00118
aldehyde metabolic process 0.00335
carbohydrate transmembrane transport 0.00588
intracellular glucose homeostasis 0.0114
glucan biosynthetic process 0.0152
mannan biosynthetic process 0.0163
trehalose biosynthetic process 0.0164
symbiont entry into host 0.0164
migration in host 0.0165
filamentous growth of a population of unicellular organisms in response to starvation 0.0193
cellular detoxification 0.0271
cellular response to oxidative stress 0.0285
glycerol metabolic process 0.0344
D-xylose metabolic process 0.0346
response to biotic stimulus 0.0384
ceramide biosynthetic process 0.047
fungal-type cell wall (1->3)-beta-D-glucan metabolic process 0.0473
Adjusted P value
C
Biological Processes for Downregulated Genes in ire1DX
endonucleolytic cleavage in ITS1 to separate SSU-rRNA from 5.8S rRNA and LSU-rRNA from tricistronic rRNA transcript (SSU-rRNA, 5.8S rRNA, LSU-rRNA) 1.6e-08
endonucleolytic cleavage to generate mature 5'-end of SSU-rRNA from (SSU-rRNA, 5.8S rRNA, LSU-rRNA) 6.62e-08
endonucleolytic cleavage in 5'-ETS of tricistronic rRNA transcript (SSU-rRNA, 5.8S rRNA, LSU-rRNA) 6.81e-08
tricarboxylic acid cycle 3.95e-06
cellular response to xenobiotic stimulus 9.61e-06
ribosomal large subunit export from nucleus 0.000284
dicarboxylic acid metabolic process 0.000315
protein import into mitochondrial matrix 0.000415
transcription initiation at RNA polymerase I promoter 0.000459
tRNA modification 0.00114
assembly of large subunit precursor of preribosome 0.00141
purine nucleobase biosynthetic process 0.00189
positive regulation of transcription by RNA polymerase I 0.00466
fluconazole transport 0.00765
CTP biosynthetic process 0.00772
purine ribonucleoside triphosphate biosynthetic process 0.00818
termination of RNA polymerase I transcription 0.00834
proton transmembrane transport 0.012
mitochondrial translation 0.0146
rRNA methylation 0.0172
mitochondrial transport 0.0184
'de novo' IMP biosynthetic process 0.0208
adenine metabolic process 0.0209
isocitrate metabolic process 0.0229
mitochondrial large ribosomal subunit assembly 0.0232
endonucleolytic cleavage in ITS1 upstream of 5.8S rRNA from tricistronic rRNA transcript (SSU-rRNA, 5.8S rRNA, LSU-rRNA) 0.0234
transcription elongation by RNA polymerase I 0.0242
protein insertion into mitochondrial inner membrane 0.0244
transition metal ion transport 0.0301
pyrimidine nucleobase biosynthetic process 0.031
xenobiotic detoxification by transmembrane export across the plasma membrane 0.0319
phosphate ion transport 0.0465
mitochondrial transcription 0.0466
Adjusted P value
D
Biological Processes for Upregulated Genes in Scire1ΔLD
cellular carbohydrate metabolic process 6.6e-06
carbohydrate metabolic process 5.06e-05
glycogen metabolic process 0.000384
energy reserve metabolic process 0.000527
cellular carbohydrate biosynthetic process 0.00222
glycogen biosynthetic process 0.00261
cellular glucan metabolic process 0.00277
glucan metabolic process 0.00277
cellular polysaccharide metabolic process 0.00342
polysaccharide metabolic process 0.00729
carbohydrate catabolic process 0.00782
glucan biosynthetic process 0.0193
cellular polysaccharide biosynthetic process 0.0217
polysaccharide biosynthetic process 0.0243
carbohydrate biosynthetic process 0.0459
transferase activity, transferring hexosyl groups 0.0486
P value
E
Biological Processes for Downregulated Genes in Scire1ΔLD
cellular amino acid metabolic process 2.83e-10
carboxylic acid metabolic process 1.22e-09
small molecule biosynthetic process 1.79e-09
oxoacid metabolic process 2.31e-09
organic acid metabolic process 2.5e-09
cellular amino acid biosynthetic process 2.31e-08
alpha-amino acid metabolic process 3.9e-08
carboxylic acid biosynthetic process 7.75e-08
organic acid biosynthetic process 7.75e-08
alpha-amino acid biosynthetic process 2.69e-07
small molecule metabolic process 4.86e-07
arginine biosynthetic process 0.000265
glutamine family amino acid metabolic process 0.00045
aromatic amino acid family metabolic process 0.000697
arginine metabolic process 0.00111
aromatic amino acid family biosynthetic process 0.00378
glutamine family amino acid biosynthetic process 0.0109
P value

**Supplemental Figure 3. RNA sequencing gene ontology analyses. A.** Principal component (PC) analysis of control (*ire1DX+IRE1-WT*) and *ire1ΔLD* grown untreated for four hours. Each strain was prepared in quadruplicate. Gene ontology analysis for biological processes was performed for *C. albicans* genes **B.** upregulated (n=675) or **C.** downregulated (n=757) in untreated *ire1DX* relative to control (Stack-Couture et al. 2026). Analysis for B and C was performed using PANTHER via the Gene Ontology Consortium (Ashburner et al. 2000; Gene Ontology Consortium 2026; Thomas et al. 2022). Gene ontology analysis for biological processes in *S. cerevisiae* genes **D.** upregulated (n=16) or **E.** downregulated (n=22) in untreated *Scire1ΔLD* relative to the control. Analysis for D and E was performed using SPELL (Version 2.0.3r71) (Hibbs et al. 2007). Terms with p value<0.05 are shown.
